# HIV-1 infection causes depletion of monocytic cells through a non-canonical cell death pathway

**DOI:** 10.1101/2022.12.20.521186

**Authors:** Michelle Stuck, Nicolas D. Arnow, Julia Kazmierski, Felix Walper, Christine Goffinet, Norbert Bannert, Oya Cingöz

## Abstract

Programmed cell death is a regulatory mechanism to eliminate infected or damaged cells. Several programmed cell death pathways exist, including apoptosis, necroptosis and pyroptosis, which can be distinguished by the cellular molecules involved. Here we show that infection of monocytic cells with HIV-1 causes cell death, which is dose- and cell type dependent and occurs independently of nucleic acid sensing or interferon (IFN) signaling. Death is observed in the case of near full-length viruses that produce viral proteins upon infection, but not in case of a minimal lentiviral vector that does not express viral gene products, demonstrating the necessity of viral gene products or a near-full length RNA genome to trigger death. Inhibition of reverse transcription or integration rescues cells, indicating that a step after integration is responsible. Using mutant viruses, we further narrow down the step in the retroviral replication cycle that triggers death. Inhibition of diverse cell death pathways individually cannot rescue cells from death following infection, consistent with PANoptosis, a cellular death process that cannot be accounted for by any single programmed cell death pathway alone. Our results elucidate the viral and cellular determinants of cell death caused by HIV-1 infection and outline cellular responses that result in the depletion of specific cell populations.

## Introduction

Programmed cell death is an essential regulatory mechanism to limit the spread and persistence of viral infections, thereby serving as a major component of intrinsic immune defense. Consequently, cell death is commonly observed during viral pathogenesis and multiple viruses have evolved strategies to manipulate specific cellular death pathways for their benefit (reviewed in [1]). Acute, untreated HIV-1 infection results in the progressive loss of CD4+ T-cells accompanied by exhausted immune compartments leading to the irreversible, immunosuppressed stage of infection defined as AIDS. Multiple programmed cell death pathways have been considered to contribute to viral pathogenesis in general, each of which can be distinguished from one another by virtue of the cellular players involved and the induced immunological response [2, 3]. Among the programmed cell death pathways apoptosis, necroptosis and pyroptosis are the most commonly observed ones in HIV-1-mediated depletion of specific cellular subsets [4-7].

Apoptosis is triggered by the presence of a pro-apoptotic stimulus resulting in the characteristic fragmentation of DNA and membrane blebbing. Apoptotic cells are eventually cleared through phagocytosis without alerting surrounding immunity, dubbing it the “cellular suicide” among the programmed death pathways [8]. Necroptosis, in contrast, is the consequence of death receptor or pattern recognition receptor (PRR)-mediated activation of the RIP3 kinase signaling pathway [9, 10]. RIP3 kinase signaling triggers cell rounding and swelling before the irreversible bursting of plasma membranes, through which immune-stimulatory cytokines and DAMPs are released to recruit immune cells and inflammation [11, 12]. Pyroptosis is an inflammatory cell death, requiring caspase-dependent signaling and installation of the inflammasome, a multimeric complex catalyzing the production of inflammation-driving cytokines, like IL-1β and IL-18, and the pore forming protein Gasdermin D to initiate inflammation [13]. Although pyroptosis, apoptosis and necroptosis harbor distinctive features, highly coordinated crosstalk between the three pathways had been observed and been accounted for by the recent implementation of the term PANoptosis [14, 15].

The characteristics of cellular death occurring upon HIV-1 infection are highly dependent on the host cell type, viral strain and experimental conditions studied. For example, CD4+ T-cells derived from lymphoid tissue undergo pyroptosis following the sensing of accumulated reverse transcription (RT) products in the cytosol caused by abortive infection, while productive infection of blood-derived CD4+ T-cells triggers apoptosis [6, 16-18]. Cells of the myeloid lineage, such as macrophages or monocyte-derived macrophages, however, largely resists the induction of cellular death pathways while exhibiting continuous virus production [19-21].

We previously reported the occurrence of cell death in THP-1 cells inoculated with high virus inputs accompanied by the induction of an antiviral IFN-stimulated gene (ISG) response [22]. In this study, we further characterize the cellular and viral determinants of this virus-induced cell death in monocytic cells upon HIV infection.

## Results

### HIV-1 infection causes death of THP-1 cells

Our lab previously showed that the triggering of ISG responses by HIV-1 reporter viruses in THP-1 cells requires high virus inputs (∼1 µg/ml p24), which simultaneously results in high levels of cell death. [22] To explore the cause of death further, THP-1 reporter cells carrying an ISRE-driven luciferase reporter (THP-1 Dual cells; Invivogen) were infected with serial dilutions of a single-round, VSV-G-pseudotyped NL43-based GFP-expressing virus (NL43-GFP). Cell death was observed in a dose-dependent manner, and at higher virus inputs, the GFP levels started to decrease concomitant with the higher level of cell death observed (Fig 1A). Treatment with nevirapine (NVP) or raltegravir (RLT) both rescued cell death, indicating a requirement for reverse transcription and integration (Fig. 1B).

**Figure 1.**
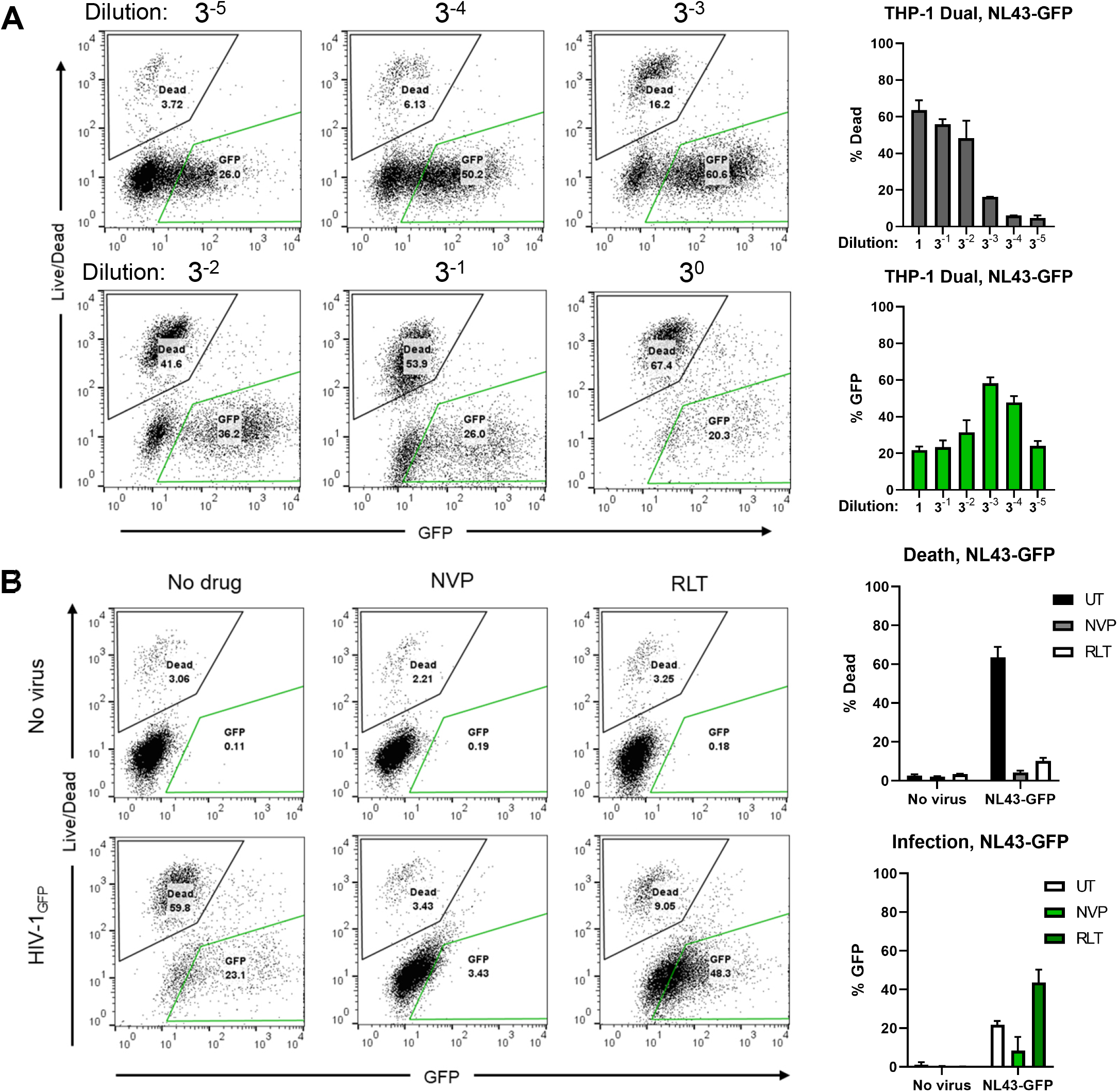
Infection of undifferentiated THP-1 cells with NL43-GFP causes cell death. (A) THP-1 Dual cells were infected with VSV-G pseudotyped, single-cycle NL43-GFP (HIV-1-GFP) virus with three-fold serial dilutions (starting concentration: 1 µg/ml p24). Cell death and infection were scored by flow cytometry two days later using live/dead staining and GFP expression and quantified. (B) THP-1 Dual cells were infected with NL43-GFP in the presence or absence of reverse transcriptase and integrase inhibitors (NVP and RLT; 10 µM each). Infection and cell death were scored as in (A). UT: untreated (no drug), NVP: nevirapine, RLT: raltegravir.

To ensure that the observed cell death was not observed in a specific clone of THP-1 cells, we tested several THP-1 clones, as well as the reporter cell line THP-1 Lucia ISG (Invivogen) in their undifferentiated state. When cells were infected with NL43-GFP, we observed a dose-dependent increase in infection levels and cell death in all cases (Fig. S1A-B), as well as type I IFN production, which was measured by incubating supernatants from infected cells on HEK-Blue IFN-α/β SEAP reporter cells (Fig S1C). Inhibition of reverse-transcription abolished all three processes, whereas inhibition of integration reduced GFP expression and partially rescued the cell death, although had no effect on ISRE induction (Fig. S1A-C). Flow cytometry profiles revealed distinct populations of dead cells and GFP-expressing cells (Fig. 1A-B) consistent with prior findings [22]. These results indicate that i) high levels of virus input are necessary to stimulate IFN responses in THP-1 cells, ii) cell death and type I IFN production in response to infection requires reverse transcription and integration, and iii) it is not the productively infected, GFP-expressing cells that seem to be dying, reminiscent of the death of CD4+ T-cells observed during abortive infection [3], although it is possible that productively infected cells would die before reaching a significant GFP expression level.

### Cell death is not caused by the activation of the cGAS/STING pathway

Pathogen sensing can cause inflammasome activation followed by pyroptosis. In human myeloid cells, the cGAS/STING axis functions upstream of the NLRP3 inflammasome [23] and it was previously reported that retroviral RT products can be sensed by the cGAS/STING pathway [24-26]. We therefore looked into whether STING KO cells, which are defective in cGAS-mediated cytoplasmic DNA sensing pathway, are rescued from virus-induced death. Infection with NL43-GFP yielded comparable levels of GFP expression and cell death in both cell types (Fig. 2A-B), indicating that STING-mediated activation of inflammasome pathway is not responsible. Virus-induced ISRE reporter expression was diminished in STING KO cells (Fig. 2C), consistent with the sensing of reverse transcription products in undifferentiated THP-1 cells. Notably, the basal level of ISRE signaling was lower in STING KO compared to WT cells, whereas the basal level of cell death was increased. RT inhibition successfully blocked infection, death and ISRE induction. While integrase inhibition reduced GFP expression and had no effect on ISRE induction as previously observed, it did rescue the cells from dying, demonstrating that cell death occurs independently of the recognition of RT products by the cGAS/STING axis.

**Figure 2.**
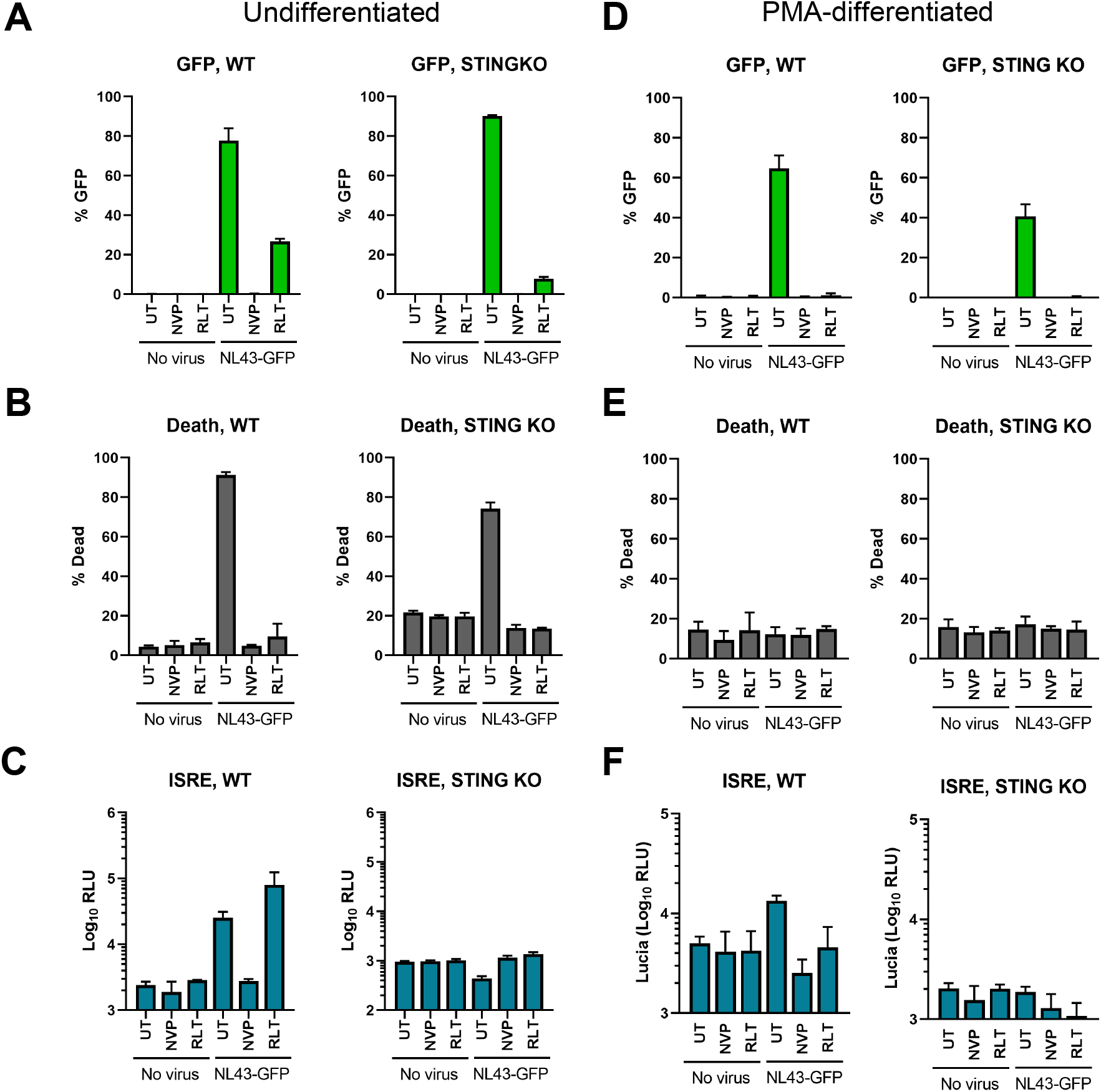
Cell death occurs independent of cGAS/STING signaling and PMA-differentiation prevents cell death. (A-C) THP-1 Dual WT and STING KO cells were infected with NL43-GFP or left uninfected, in the presence or absence of RT and IN inhibitors. Infection (A) and death (B) were quantified by flow cytometry. ISRE induction (C) was quantified by a luciferase assay from the supernatants of infected cells. (D-F) Experiments were performed exactly as in (C-E), but in overnight PMA-differentiated (25 ng/ml; 24h) THP-1 Dual WT and STING KO cells. ISRE: IFN-stimulated response element.

### PMA-differentiation of THP-1 cells rescues infection-induced cell death

Although not an ideal system, THP-1 cells differentiated by PMA are widely used to mimic macrophage-like cells in tissue culture. To test whether the differentiation status had an effect on virus-induced cell death, we performed the same experiments in Fig. 2A-C using PMA-differentiated THP-1 Dual WT and STING KO cells to quantify infection, cell death and ISRE reporter expression. Infection levels in differentiated cells were generally lower than their undifferentiated counterparts (Fig 2A vs. D), although comparable between WT and STING KO cells. Interestingly, both basal and infection-induced cell death was completely blocked by PMA treatment (Fig. 2E). Fold ISRE expression due to infection was reduced in PMA-differentiated cells compared to undifferentiated cells (Fig. 2C vs. F), which could be due to the lower infection levels or the altered ability to respond to different stimuli upon differentiation. These results demonstrate that PMA-differentiation rescues cells from dying upon infection.

### SAMHD1 deficiency does not rescue cell death caused by infection

It was reported that SAMHD1 controls the cell cycle and that its deletion results in decreased sensitivity to apoptosis in THP-1 cells [27]. We therefore checked whether lack of SAMHD1 would rescue infection-induced cell death. We infected two different clones of SAMHD1 KO THP-1 cells (Fig. S2A) with increasing doses of NL43-GFP in the presence or absence of NVP or RLT. As in WT cells, we also observed a dose-dependent increase in GFP expression and cell death, although the two clones showed some variation in terms of their sensitivity to death (Fig. S2B-C). For SAMHD1 KO clone #1, we also observed a dose-dependent increase in type I IFN production, whereas clone #2 did not have any detectable type IFN (Fig. S2D). At higher doses of virus, this lack of response could be due to the high level of cell death (∼80%), although at lower doses it is likely due to clonal differences between the two lines. Overall, these results show that lack of SAMHD1 does not influence cell death caused by NL43-GFP infection.

### Exposure to virus, not a secreted cellular factor, mediates cell death

We observed that productively infected cells (i.e. expressing GFP) and dead cells formed two distinct populations (Fig. 1A), which could indicate bystander cell death. To determine whether a secreted factor was responsible, we performed supernatant removal and transfer experiments. After removing the inoculated virus, supernatants from infected THP-1 cells in the presence or absence of NVP were collected at 24 and 48 hours, and added to naïve THP-1 cells. Two days of incubation with conditioned supernatants did not cause an increase in cell death over basal levels, and neither did the inhibition of RT (Fig. 3A). Conversely, the removal of the media after infection to prevent continuous exposure to potential danger-associated factors released from cells failed to rescue death in cells that had been exposed to the virus (Fig. 3B). These results indicate that death is not caused by a secreted factor and that exposure to the virus is necessary, although we cannot distinguish whether it is the abortively infected cells or bystander cells that are dying in this case.

**Figure 3.**
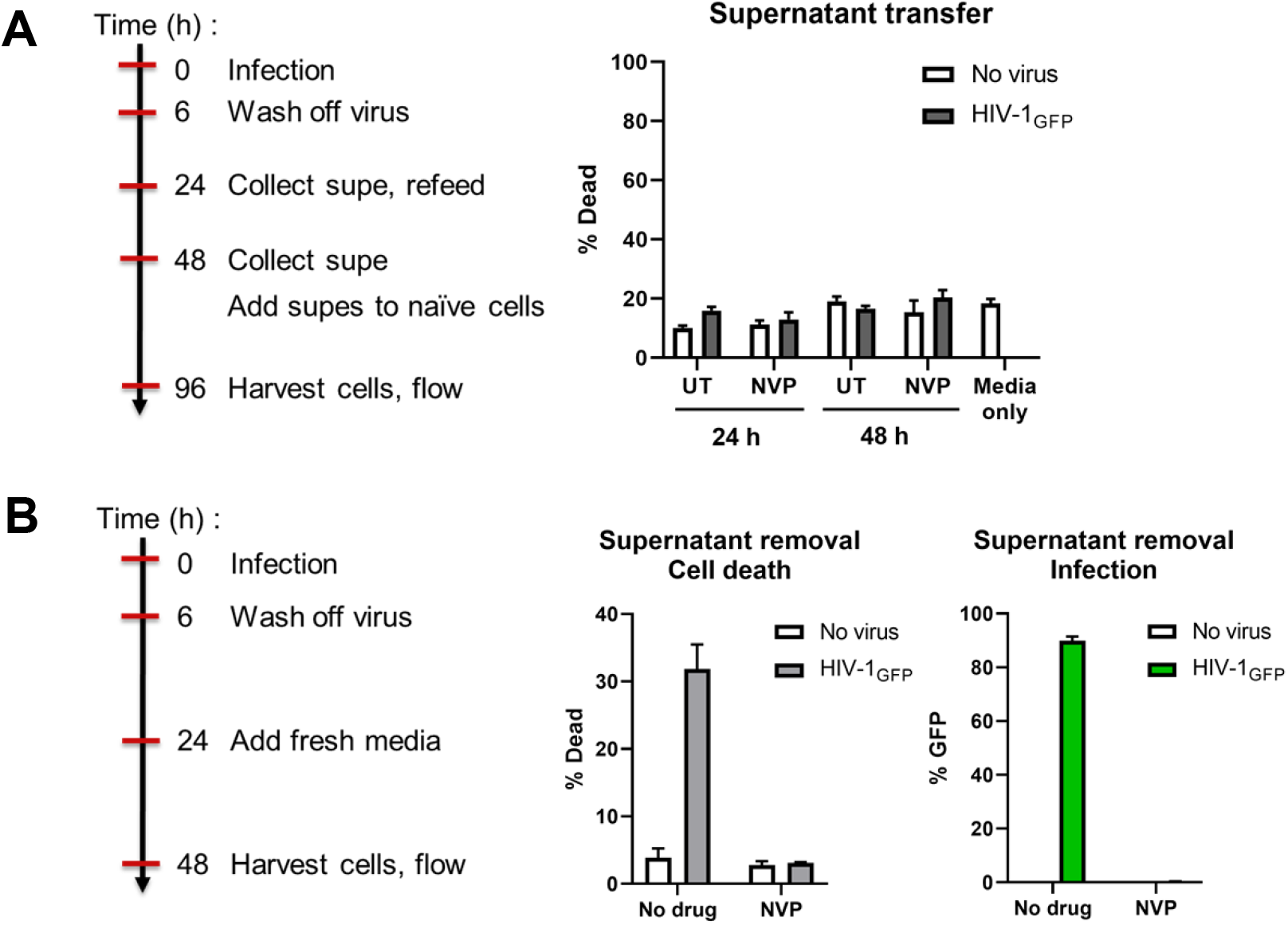
Cell death requires exposure to the virus and is not due to a secreted factor. (A) THP-1 WT cells were infected with NL43-GFP or not, in the presence or absence of NVP (10 µM). After removal of the virus, conditioned supernatants were collected at 24 and 48 hours and added to naïve cells. Cell death was assessed two days later by flow cytometry. (B) THP-1 WT cells were infected or not with NL43-GFP, in the presence or absence of NVP (10 µM). Media was replaced after 6 and 24 hours. Cell death and infection were assessed by flow cytometry 48 hpi.

### Death due to HIV-1 infection is cell-type dependent

We next tested whether the death of cells following HIV-1 infection is a common feature of monocytic cell lines. Similar to our observations in THP-1 cells, infection of MonoMac-6 cells with HIV-1-GFP caused a dose-dependent increase in death, whereas U937 cells were largely resistant (Fig. 4A-B). Infection of primary monocytes isolated from PBMCs of healthy donors with HIV-1-GFP and HIV-1-Luc viruses in the presence of Vpx-containing VLPs also did not result in higher levels of cell death than controls (Fig. 4C), demonstrating that cellular demise after HIV-1 infection is cell-type dependent. It should be mentioned that following infection, primary monocytes showed morphological changes that suggested their differentiation into macrophage-like cells.

**Figure 4.**
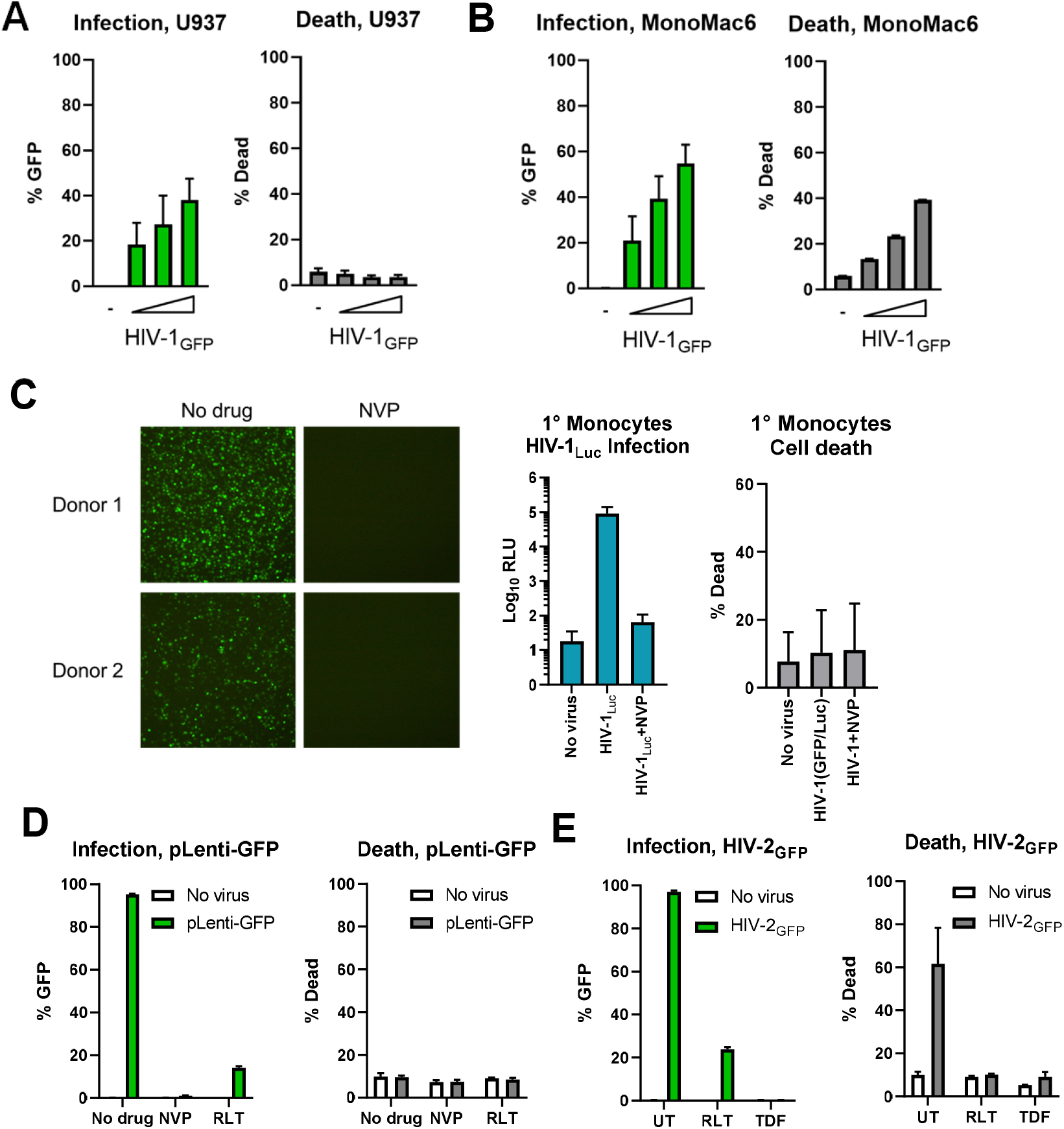
Infection-induced death is cell type dependent and requires the presence of viral genes. (A-B) U937 or MonoMac6 monocytic cell lines were infected with NL43-GFP in the presence or absence of NVP (10 µM). Cell death and infection rates were analyzed by flow cytometry. (C) Primary monocytes from healthy donors were infected with NL43-GFP(+Vpx) or NL43-Luc(+Vpx). Infection was assessed by microscopy and by firefly luciferase assay, respectively, while cell death was assessed by flow cytometry. (D) THP-1 cells were infected with a minimal lentiviral vector, pLenti-GFP, in the presence or absence of NVP or RLT (10 µM). Infection and death were measured as in (A). (E) Similar to (D), except cells were infected with VSV-G pseudotyped, ROD-10 based, single-round HIV-2.

### The presence of viral genes or a near full-length genome is necessary for infection-mediated cell death

To delineate whether the presence of one or more viral genes was mediating the cell killing during infection, we employed a minimal lentiviral vector (pLenti-GFP), which can reverse transcribe and integrate, but does not encode any of the lentiviral genes. Infection of THP-1 cells VSV-G pseudotyped pLenti-GFP resulted in high infection levels, but did not induce death (Fig. 4D), indicating that the presence of a near full-length viral genome or viral protein expression is necessary for death. In agreement with these results, infection with a VSV-G-pseudotyped, ROD10-based HIV-2_GFP_ reporter virus behaved similar to HIV-1, resulting in cell death, whereas inhibition of reverse transcription and integration prevented death (Fig. 4E).

### Gag expression alone does not cause cell death

We reasoned that a high level of Gag expression and continuous budding of VLPs from the cell membrane might overwhelm the cells and compromise the integrity of the plasma membrane. To test whether cell death is due to Gag expression, we created a tet-inducible THP-1 cell line that expresses a codon-optimized HIV-1 Gag upon stimulation. Doxycycline treatment resulted in a ∼55-60% Gag expression at 1-2 days post-infection (Fig. 5A), concomitant with particle release into culture supernatants (Fig. 5B). We did not observe a marked increase in cell death due to infection; although the levels were slightly elevated at 48 h, they remained well below 10%, suggesting that Gag expression alone does not recapitulate the death caused by infection.

**Figure 5.**
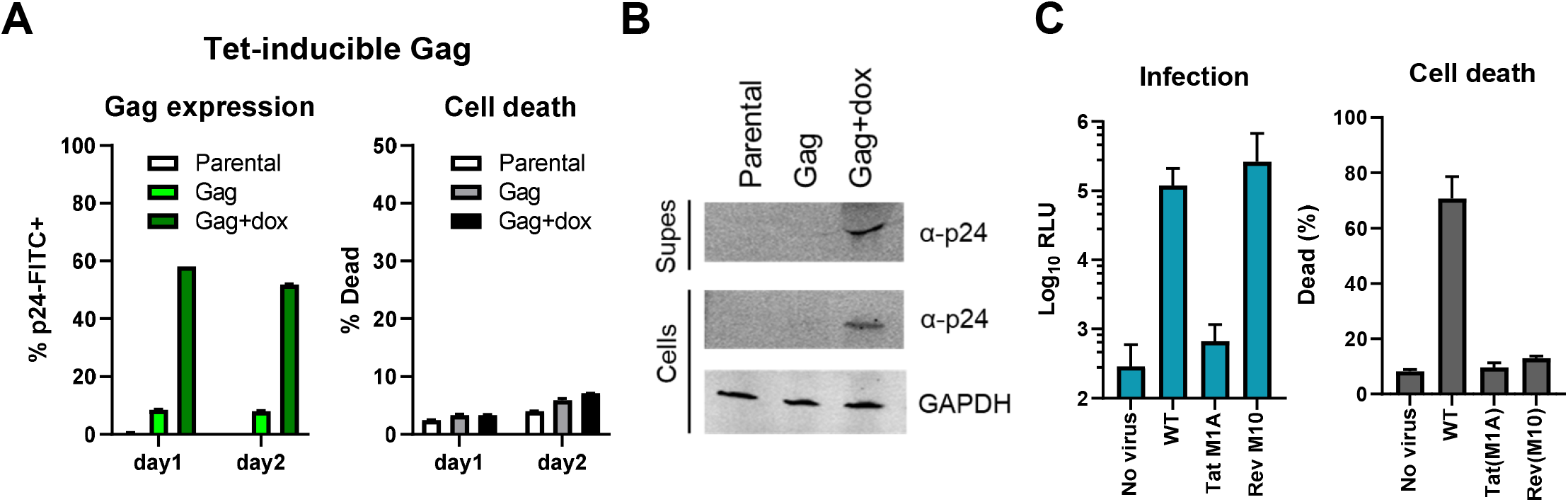
Death occurs at a step after the nuclear export of intron-containing viral RNA, but is not due to Gag overexpression. (A) Tet-inducible (Tet-ON) THP-1 cells were stably transduced with a codon-optimized Gag expressing vector. Cells were stimulated with doxycycline (dox; 1 µg/ml) and Gag expression was quantified by p24 staining at 24 and 48 h after treatment. (B) As in (A), except cells and supernatants were harvested, supernatants were centrifuged to pellet virus like particles, and the resulting lysates were probed for p24 expression on a Western blot. (C) THP-1 cells were infected with Tat or Rev mutant NL43-Luc viruses. Infection and death were quantified by firefly luciferase assay and live/dead staining, respectively.

The requirement for reverse transcription and integration pointed to a post-integration step that is responsible for cell death. To determine which steps are necessary, we used NL43-based mutants that lack functional tat or rev genes. The Tat mutant has an alanine instead of the start codon (M1A) and cannot transcribe efficiently from the provirus, whereas the rev mutant (M10; LE76/77DL) is a well-described dominant-negative version that cannot export intron-containing viral RNAs from the nucleus to the cytoplasm for translation or packaging [28]. Infection levels with the Tat mutant was similar to uninfected cells, which was expected as proviral transcription is necessary for reporter expression. Infection with the Rev mutant virus was similar to WT, as the firefly luciferase reporter gene in this construct is produced from multiply-spliced viral RNA and does not require Rev for expression. Notably, the absence of functional Tat or Rev proteins rescued infected cells from dying (Fig. 5C), illustrating that the step at which cell death occurs takes place after the nuclear export of RRE-containing viral RNAs. These data also suggest that either the presence of intron-containing (or near full-length) viral RNA in the cytoplasm, or expression of viral proteins from such RNA species may be required for triggering cell death during infection.

### HIV-1 infection causes IL-1β, but not IL-18, mRNA expression

Pyroptosis, a highly inflammatory form of cell death, is regulated by cellular caspase activity. Activation of Caspase 1 during pyroptosis results in the proteolytic processing of pro-inflammatory cytokines pro-IL-1β and pro-IL-18, converting them to their active forms, IL-1β and IL-18, which are then released from dying cells. Based on reports that HIV-1 infection can activate pyroptosis under certain conditions [6, 29], we reasoned that this might explain the cell death observed in our experimental setting. Infection of undifferentiated THP-1 cells with NL43-GFP induced a significant increase (>four-log) in IL-1β mRNA expression at 24 h, whereas IL-18 mRNA levels did not change (Fig. 6A). NVP treatment or PMA-differentiation of cells prevented the virus-mediated increase in IL-1β mRNA. The stimulation of IL-1β expression was dose-dependent, which mirrored the expression of several other ISGs, including CXCL10, IFIT1 and IFIT2 (Fig. 6B). These results suggested that the cellular death pathway might be linked to pyroptosis, although no IL-1β secretion could be detected in culture supernatants of infected cells (data not shown).

**Figure 6.**
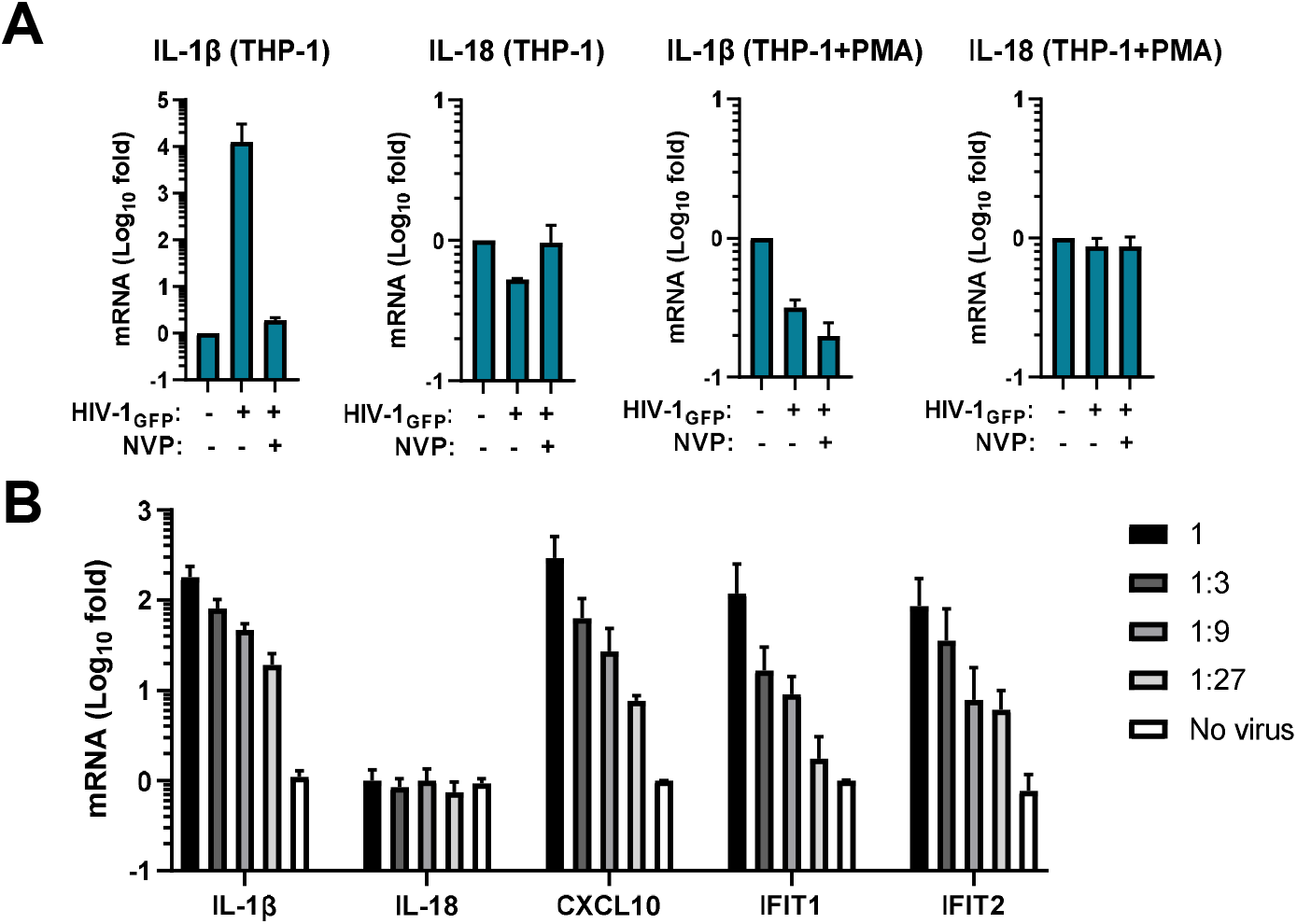
HIV-1 infection stimulates IL-1β, but not IL-18 expression. (A) Undifferentiated or PMA-differentiated THP-1 cells were infected with NL43-GFP (HIV-1-GFP). IL-1β and IL-18 mRNA levels were quantified by RT-qPCR two days after infection. (B) Undifferentiated THP-1 cells were infected with three-fold serial dilutions of NL4.3-GFP. mRNA levels for the indicated genes were measured by RT-qPCR two days after infection.

### Inhibitors of various cellular death pathways cannot rescue infection-induced death

HIV-1 infection can activate various nucleic acid sensing and inflammasome pathways, through cGAS/STING, MAVS, TLRs, DDX3, IFI16, and CARD8, among others [24, 26, 29-35]. Depending on the cell type, extracellular environment and the viral strains used, HIV-1 infection has been reported to cause cell death via apoptosis, necroptosis or pyroptosis [3-5, 7]. Autophagy-inducing peptides can selectively kill HIV-1 infected cells in a caspase-independent form of cell death called autosis [36]. HIV-1 was also shown to cause CD4+ T-cell death due to integration, in a DNA-PK-dependent manner [37]. To elucidate whether any of these pathways may be involved in our system, we used chemical inhibition of various host targets and assessed cell death after infection. Inhibition of IFN signaling by ruxolitinib, DNA-PK activity by AZD-7648, endosomal acidification and autophagy by bafilomycin A and chloroquine, or autosis using digoxin [38] did not relieve virus-induced cell death, while none of the drugs were toxic at the given concentrations in the absence of virus (Fig. 7A). The only drugs to prevent death were the two antiretrovirals, RT and IN inhibitors, as mentioned (Fig. 1D). To look more closely into the various cell death pathways, treatment of cells with a pan-caspase inhibitor (Z-VAD-FMK) to block apoptosis and pyroptosis, a caspase 1-specific inhibitor (VX-765) to block pyroptosis, or inhibitors of RIPK1 and MLKL (necrostatin and necrosulfonamide, respectively) to block necroptosis failed to rescue virus-induced cell death (Fig. 7B). None of the drugs were toxic in the absence of virus, with the exception of necrosulfonamide that showed a dose-dependent increase in cell death even in the absence of infection. Our results show that inhibition of the cell death pathways tested here cannot rescue HIV-1 infected monocytic cells from death.

**Figure 7.**
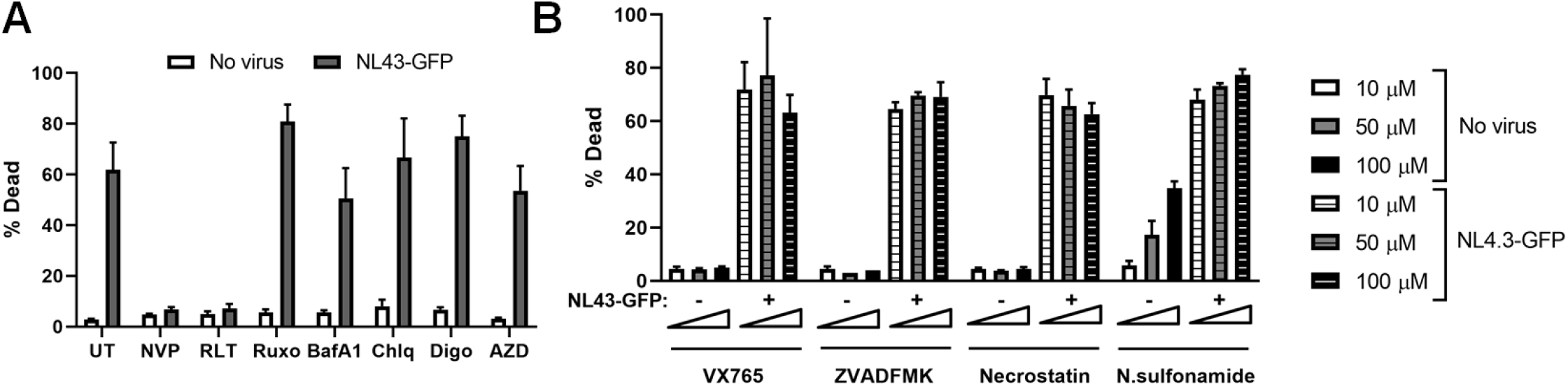
Inhibition of autophagy, apoptosis, pyroptosis, necroptosis or autosis do not rescue cell death caused by HIV-1 infection. (A) THP-1 cells were treated with the inhibitors nevirapine (NVP; 10 µM), raltegravir (RLT; 10 µM), ruxolitinib (Ruxo; 1 µM), bafilomycin A1 (BafA1; 10 µM), chloroquine (Chlq; 10 µM), digoxin (Digo; 0.1 µM) or AZD-7648 (AZD; 10 µM) at the time of infection with NL43-GFP. Cell death was quantified two days after infection. (B) THP-1 cells were treated with inhibitors of apoptosis (Z-VAD-FMK, pan-caspase inhibitor), pyroptosis (VX765, caspase 1 inhibitor) and necroptosis (necrostatin, RIPK1 inhibitor and necrosulfonamide, MLKL inhibitor) at the indicated concentrations. Cells were challenged with NL43-GFP or mock infected, and the level of cell death was measured after two days.

## Discussion

We show here that infection of monocytic cells in culture with a near-full length HIV-1 triggers cell death. The phenotype we observed was cell type dependent, as some monocytic cells such as THP-1 and MonoMac-6 were susceptible, whereas others such as U937 were largely resistant. Differentiation of cells prior to infection reversed this susceptibility and conferred resistance against death by infection. Cell death was observed with HIV-1 as well as HIV-2-based viruses, which could be prevented by inhibiting reverse transcription or integration by antiretroviral drugs, suggesting that a step after integration is responsible. Blocking proviral transcription or nuclear export of intron-containing viral RNAs also prevented death. In alignment with these findings, infection with minimal lentiviral vectors expressing GFP did not phenocopy the death observed with near full-length HIV-1. Thus, we conclude that the expression of one or more viral genes or the presence of intron-containing viral RNA forms in the cytoplasm is necessary for HIV-induced cell death.

It remains to be seen which exact viral determinant is responsible for cell death we observe during infection. Expression of Gag alone, in the absence of any other viral genes, is typically sufficient to form virus-like particles (VLPs)[39]. We reasoned that high levels of infection could lead to constant Gag production and budding of VLPs, which could compromise the integrity of the plasma membrane. Overexpression of a codon-optimized HIV-1 Gag alone, however, did not lead to cell death, despite reaching high expression levels and VLP production in culture supernatants (Fig. 5A-B). The auxiliary protein Vpr induces cell cycle arrest followed by apoptosis [40], the Env glycoprotein can induce bystander cell apoptosis [41] and the Nef protein can cause endothelial cell death [42]; however, the NL43-based vector used in our study is deficient for all three genes. Overexpression of Rev in cultured cell lines such as 293T and HeLa was reported to lead to cell death, which was only observed in nondividing cells [43]. We therefore believe that Rev accumulation also does not explain the death observed in our experiments, as our cells are dividing, and PMA differentiation, which pushes them to a nondividing phenotype, had the opposite effect of rescuing cells from infection-induced death.

That primary monocytes did not show higher rates of cell death due to HIV-1 infection *ex vivo*, could raise the argument that our experimental system may not accurately reflect the conditions observed *in vivo*. It should be mentioned that in our experiments, infection of primary monocytes isolated from PBMCs with HIV-1 resulted in the differentiation of these cells into macrophage-like cells, even in the absence of additional growth factors. This MDM type differentiation was readily observed by morphological changes, which is in line with other reports that HIV-1 infection induces the activation of monocyte-derived macrophages and dendritic cells [25, 30, 31]. As undifferentiated THP-1 cells were highly susceptible to infection-induced cell death, whereas PMA-differentiation rendered them entirely resistant, it is perhaps not surprising that infection-induced differentiation of monocytes confers protection against death at the same time. This makes it quite difficult to address whether primary monocytes would phenocopy monocytic cell lines in terms of infection-induced death, unless their differentiation into MDMs could be prevented.

Our efforts to identify which cellular death pathway is activated by HIV-1 was hampered by the fact that none of the inhibitors of apoptosis, pyroptosis, necroptosis, autosis or autophagy could rescue cells from infection-induced death. It could be that more than one death pathway is activated, rendering a single inhibitor unable to prevent death; however, combinatorial treatment with several inhibitors also did not rescue cells (data not shown). Indeed, the term PANoptosis has been used to describe an inflammatory type of programmed cell death which displays features of pyroptosis, apoptosis and necroptosis, but cannot be explained by any single pathway alone [44]. Both RNA and DNA viruses, including herpes simplex virus (HSV) and influenza A virus (IAV) can activate multiple cell death pathways simultaneously during infection (reviewed in [45]); it is conceivable that HIV-1/2 might do the same. Inhibition of other cellular processes, for instance those reported to be involved in HIV-1 mediated T-cell killing such as DNA-PK [37], or IFN-induced JAK/STAT signaling also failed to prevent death. It is possible that when cells commit to a death pathway they reach a “point of no return” such that the process can no longer be reversed, although the presence of inhibitors against different cell death pathway components at the beginning of the infection would be expected to prevent this commitment in the first place. Further work would be required to investigate the potential involvement of other cell death pathways, such as ferroptosis or oxeiptosis [46, 47], which could be explored using genetic knockouts or chemical inhibitors in the future.

## Supplementary Figure legends

**Figure S1.**
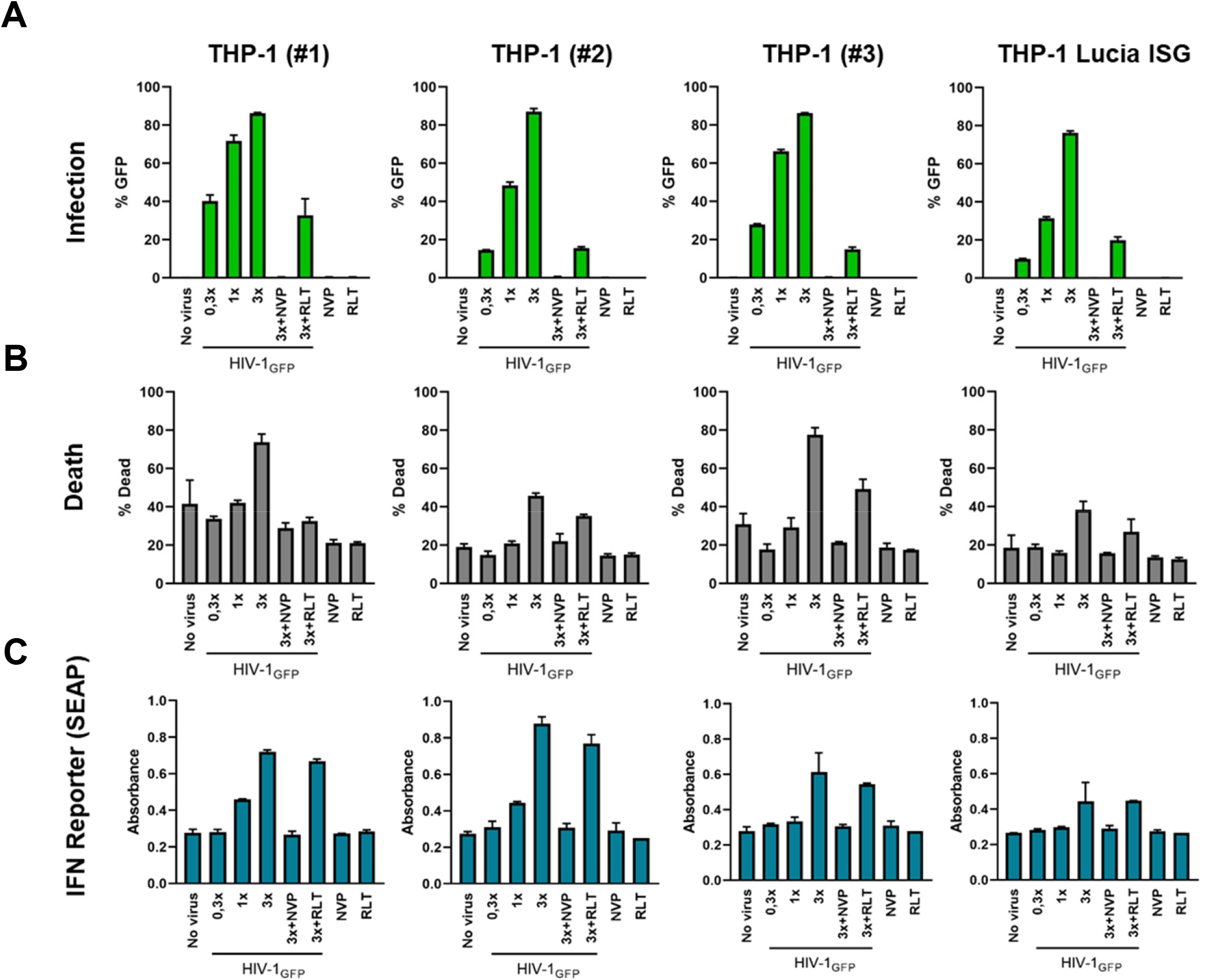
Infection, ISRE induction and cell death in various undifferentiated THP-1 clones. Undifferentiated THP-1 cells from different sources were infected with VSV-G pseudotyped, single-cycle NL43-GFP at three different viral inputs in the presence or absence of reverse transcriptase and integrase inhibitors (NVP and RLT; 10 µM). (A-B) Infection and cell death were scored by flow cytometry using GFP expression and live/dead stain. C) Supernatants from infected cells were assayed on HEK-Blue IFN-α/β SEAP reporter cells.

**Figure S2.**
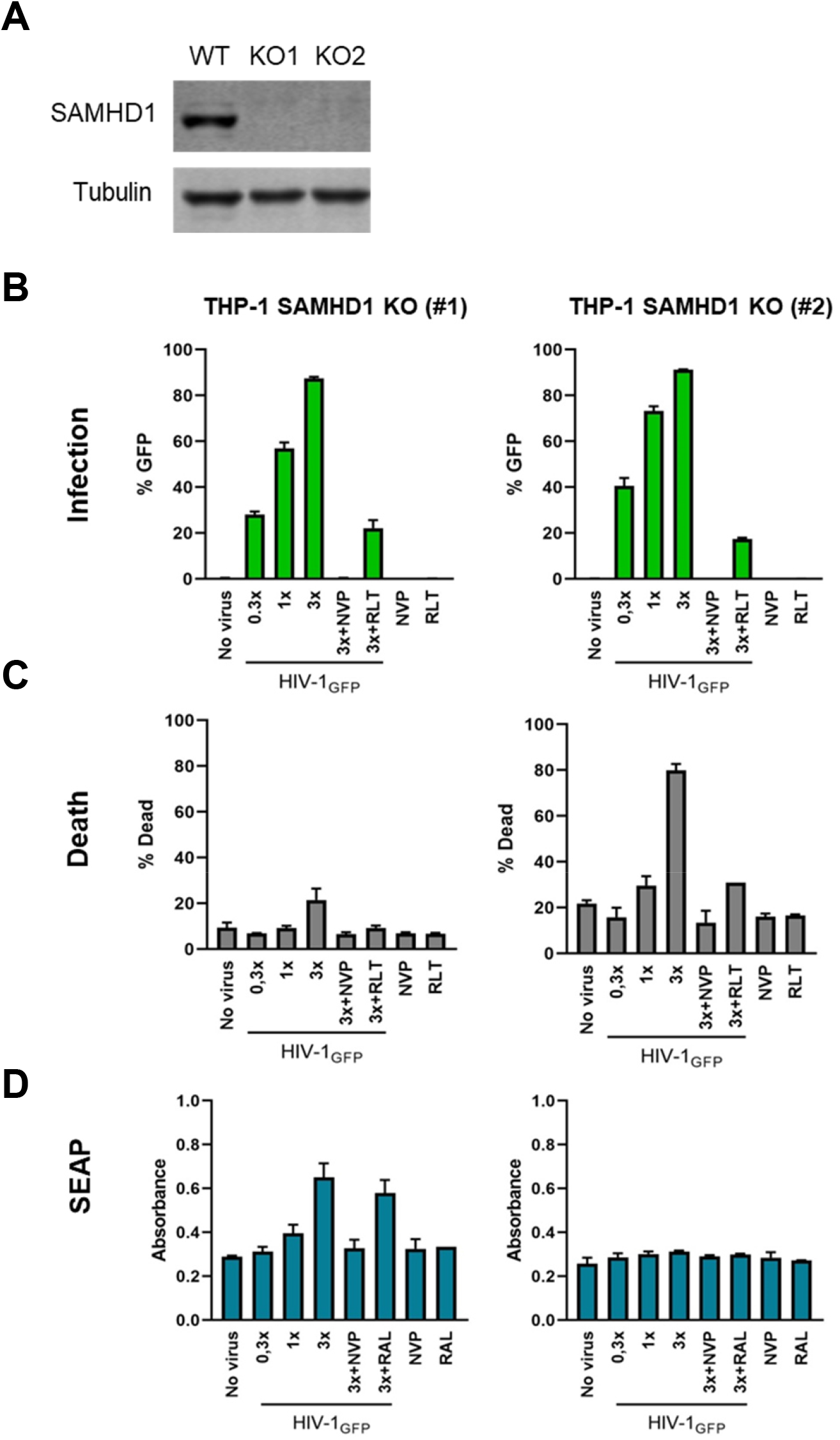
SAMHD1 deficiency does not rescue cell death caused by NL43-GFP. Undifferentiated THP-1 cells deficient in SAMHD1 were infected with VSV-G pseudotyped HIV-1-GFP at different viral inputs in the presence or absence of NVP or RLT (10 µM). (A) Cell lysates from WT or SAMHD1 KO cells were analyzed by Western blot, probed for SAMHD1 and tubulin. (B-C) Infection and cell death were scored by flow cytometry. (D) Supernatants from infected cells were assayed on HEK-Blue IFN-α/β SEAP reporter cells.

## Methods

### Cell culture

293T and HEK-Blue IFN-α/β cells (Invivogen) were maintained in DMEM (Gibco) containing 9% FBS (Gibco) and 100 µg/ml Pen/Strep (ThermoFisher). THP-1 Lucia ISG cells were obtained from Invivogen. THP-1 SAMHD1 KO and WT control cells were kindly provided by Torsten Schaller. MonoMac6 and U937 cells were kindly provided by Elisabeth Kamal (RKI, Berlin). All monocytic cell lines were maintained in RPMI with 9% FBS, 100 µg/ml Pen/Strep, 100 µg/ml Normocin (Invivogen). For differentiation, THP-1 cells were treated with PMA (25 ng/ml) for 24 hours, followed by fresh media addition. PBMCs were isolated from buffy coats from the Red Cross using standard Ficoll separation. Monocytes were selected by adhering PBMCs in RPMI containing 5% pooled human serum, 1 mM HEPES and 24 µg/ml gentamicin for several hours, followed by extensive washing to remove unbound cells.

### Plasmids, virus production and infection

HIV-1-GFP was produced by transfection of the NL43-based viral plasmid pHIV-CMV-GFP SIVp6(17-26), which lacks *env, nef* and *vpr* genes, contains a 10 amino acid stretch from SIV p6 (for efficient Vpx packaging), and GFP expression is driven by a CMV promoter. The minimal lentiviral vector pLentiGFP only contains the LTRs and cis-regulatory RNA elements and expresses GFP. Both plasmids were kindly provided by Henning Hoffman. NL43-Luc was produced from pNL4-3.Luc R-E-that also lacks *env, nef* and *vpr* genes, kindly provided by Ned Landau and obtained from the HIV Reagent Program (ARP-3418). Viruses were produced by transfecting the viral plasmids together with a plasmid encoding VSV-G env (pCMV-VSV-G, kindly provided by Bob Weinberg; Addgene #8454), and in case of the minimal lentiviral vectors, also with a plasmid encoding HIV-1 *gag, pol, tat* and *rev* (psPAX2, kindly provided by Didier Trono; Addgene #12260). For virus packaging of Vpx a plasmid encoding SIVmac239 Vpx gene was transfected simultaneously. Codon-optimized HIV-1 *gag* (pTH-HIV-1 coGag) was kindly provided by Oliver Hohn (RKI). Virus stocks were produced by standard transfection of 293T cells with polyethylenimine (PEI), followed by media change after one day and supernatant collection after two days. Virus-containing supernatants were filtered, ultracentrifuged over 20% sucrose, resuspended, aliquoted and frozen at -80°C. Infections were performed by spinoculation at 1200 x g for 1-2 h at 25°C. Infectious units were determined on TZM-Bl cells followed by X-gal staining or in case of GFP viruses on 293T cells by flow cytometry.

### Flow cytometry

For assessing cell death, cells were pelleted, washed once with PBS, stained with death markers as indicated (fixable viability dye-660, eBioscience) at room temperature for 15 min, washed twice with PBS and then fixed in IC fixation buffer for 15 min at room temperature (eBioscience). For intracellular p24 staining, fixed cells were incubated with FITC-p24 antibody (KC57; Beckman Coulter) in the presence of 0.1% Triton-X. Samples were diluted in PBS prior to data acquisition by flow cytometry (FACScalibur, BD). Further analysis of different cell populations was performed in FlowJo (v10.6.2).

### Luciferase Assays

For lucia luciferase assays, supernatants from infected or treated cells were collected and plated in an opaque/white 96-well plate. Quanti-Luc reagent (Invivogen) was used as substrate. For firefly luciferase assays, cells were lysed in passive lysis buffer, which was detected with luciferase detection reagent (Promega). Luminescence measurements were done using a LUMIstar OMEGA plate reader (BMG) with automatic injections.

### RT-qPCR

For mRNA expression analysis, total cellular RNA was isolated using RNeasy Mini Kit (Qiagen), treated with Turbo DNase and inactivation reagent (Ambion), cDNA was synthesized using Superscript III (Invitrogen), and qPCR was performed with the SensiFAST No-ROX Master Mix (Bioline), all according to manufacturer’s instructions on a Bio-Rad CFX96 qPCR machine. A list of primer sequences is provided in Table 1.

**Table 1.**
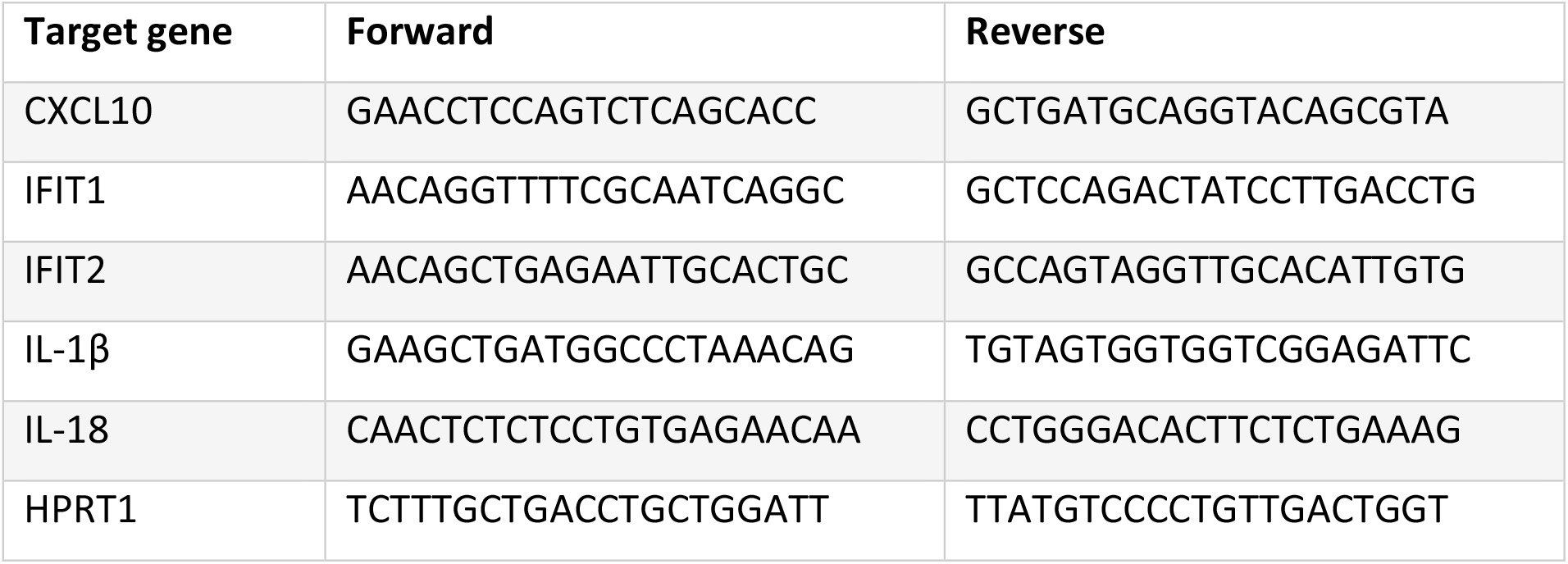
List of primers used for RT-qPCR.

### Western blot

Cells were washed with PBS and lysed in 100 mM Tris, 30 mM NaCl, 0.5% NP40 containing proteasome inhibitors (Roche). Lysates were analyzed by SDS-PAGE using standard techniques. Primary antibodies were rabbit anti-STING (CST), mouse anti-SAMHD1 (Bio-Rad), mouse anti-tubulin (Sigma); mouse anti-GAPDH (Thermo). IRdye-labeled secondary antibodies were from Licor. Blots were scanned using Odyssey infrared scanner and analyzed further using ImageStudio software (Licor).

### SEAP assay

Supernatants were collected from infected cells and incubated with freshly plated HEK-Blue IFN-α/β cells overnight in a 96-well plate (25K cells/well). The next day, supernatants from HEK-Blue cells were mixed with Quanti-Blue reagent (Invivogen) and absorbance was measured in a plate reader at 650 nm.

### Drugs

The drugs/chemicals used in this study are listed, along with the companies they were obtained from in parentheses: Nevirapine, and tenofovir (Merck), raltegravir (Santa Cruz), Ruxolitinib, AZD7648, necrosulfonamide (Selleckchem), Z-VAD-FMK, bafilomycin A1 (Invivogen), VX-765 (Tocris), MG132, chloroquine, digoxin (Sigma).

## Acknowledgements

We thank all members of the Cingöz Lab and all of Unit 18 (FG18) for helpful discussions and reagents. This work was supported by internal funds from the Robert Koch Institute (Berlin, Germany) and the German Research Foundation (DFG) Priority Program Grant: Innate sensing and restriction of retroviruses (SPP1923). We also thank the Klaus Tschira Foundation and German Scholars Organization (GSO) for the KT Boost Fund awarded to Oya Cingöz.

## References

1. Imre G. Cell death signalling in virus infection. Cellular Signalling. 2020;76:109772. doi: 10.1016/j.cellsig.2020.109772.

2. Alimonti JB, Ball TB, Fowke KR. Mechanisms of CD4+ T lymphocyte cell death in human immunodeficiency virus infection and AIDS. The Journal of General Virology. 2003;84(Pt 7):1649–61. doi: 10.1099/vir.0.19110-0.

3. Doitsh G, Greene Warner C. Dissecting How CD4 T Cells Are Lost During HIV Infection. Cell Host & Microbe. 2016;19(3):280–91. doi: https://doi.org/10.1016/j.chom.2016.02.012.

4. Pan T, Wu S, He X, Luo H, Zhang Y, Fan M, et al. Necroptosis Takes Place in Human Immunodeficiency Virus Type-1 (HIV-1)-Infected CD4+ T Lymphocytes. PloS one. 2014;9(4):e93944. doi: 10.1371/journal.pone.0093944.

5. Terahara K, Iwabuchi R, Iwaki R, Takahashi Y, Tsunetsugu-Yokota Y. Substantial induction of non-apoptotic CD4 T-cell death during the early phase of HIV-1 infection in a humanized mouse model. Microbes Infect. 2021;23(1):104767. Epub 20201010. doi: 10.1016/j.micinf.2020.10.003. PubMed PMID: 33049386.

6. Doitsh G, Galloway NL, Geng X, Yang Z, Monroe KM, Zepeda O, et al. Cell death by pyroptosis drives CD4 T-cell depletion in HIV-1 infection. Nature. 2014;505(7484):509–14. Epub 2013/12/21. doi: 10.1038/nature12940. PubMed PMID: 24356306; PubMed Central PMCID: PMCPMC4047036.

7. Garg H, Joshi A. Host and Viral Factors in HIV-Mediated Bystander Apoptosis. Viruses. 2017;9(8). Epub 20170822. doi: 10.3390/v9080237. PubMed PMID: 28829402; PubMed Central PMCID: PMCPMC5579491.

8. Singh R, Letai A, Sarosiek K. Regulation of apoptosis in health and disease: the balancing act of BCL-2 family proteins. Nature Reviews Molecular Cell Biology. 2019;20(3):175–93. doi: 10.1038/s41580-018-0089-8.

9. Holler N, Zaru R, Micheau O, Thome M, Attinger A, Valitutti S, et al. Fas triggers an alternative, caspase-8-independent cell death pathway using the kinase RIP as effector molecule. Nature immunology. 2000;1(6):489–95. doi: 10.1038/82732.

10. Silke J, Rickard JA, Gerlic M. The diverse role of RIP kinases in necroptosis and inflammation. Nature immunology. 2015;16(7):689–97. doi: 10.1038/ni.3206.

11. Pasparakis M, Vandenabeele P. Necroptosis and its role in inflammation. Nature. 2015;517(7534):311–20. doi: 10.1038/nature14191.

12. Zhou Z, Han V, Han J. New components of the necroptotic pathway. Protein & Cell. 2012;3(11):811–7. doi: 10.1007/s13238-012-2083-9.

13. Man SM, Karki R, Kanneganti T-D. Molecular mechanisms and functions of pyroptosis, inflammatory caspases and inflammasomes in infectious diseases. Immunological reviews. 2017;277(1):61–75. doi: 10.1111/imr.12534.

14. Malireddi RKS, Kesavardhana S, Kanneganti T-D. ZBP1 and TAK1: Master Regulators of NLRP3 Inflammasome/Pyroptosis, Apoptosis, and Necroptosis (PAN-optosis). Frontiers in Cellular and Infection Microbiology. 2019;9:406. doi: 10.3389/fcimb.2019.00406.

15. Samir P, Malireddi RKS, Kanneganti T-D. The PANoptosome: A Deadly Protein Complex Driving Pyroptosis, Apoptosis, and Necroptosis (PANoptosis). Frontiers in Cellular and Infection Microbiology. 2020;10:238. doi: 10.3389/fcimb.2020.00238.

16. Doitsh G, Cavrois M, Lassen KG, Zepeda O, Yang Z, Santiago ML, et al. Abortive HIV Infection Mediates CD4 T-Cell Depletion and Inflammation in Human Lymphoid Tissue. Cell. 2010;143(5):789–801. doi: 10.1016/j.cell.2010.11.001.

17. Monroe KM, Yang Z, Johnson JR, Geng X, Doitsh G, Krogan NJ, et al. IFI16 DNA Sensor Is Required for Death of Lymphoid CD4 T-cells Abortively Infected with HIV. Science (New York, NY). 2014;343(6169):428–32. doi: 10.1126/science.1243640.

18. Muñoz-Arias I, Doitsh G, Yang Z, Sowinski S, Ruelas D, Greene WC. Blood-Derived CD4 T Cells Naturally Resist Pyroptosis During Abortive HIV-1 Infection. Cell host & microbe. 2015;18(4):463–70. doi: 10.1016/j.chom.2015.09.010.

19. Bagnarelli P, Valenza A, Menzo S, Sampaolesi R, Varaldo PE, Butini L, et al. Dynamics and modulation of human immunodeficiency virus type 1 transcripts in vitro and in vivo. Journal of Virology. 1996;70(11):7603–13.

20. Cassol E, Alfano M, Biswas P, Poli G. Monocyte-derived macrophages and myeloid cell lines as targets of HIV-1 replication and persistence. Journal of Leukocyte Biology. 2006;80(5):1018–30. doi: 10.1189/jlb.0306150.

21. Igarashi T, Brown CR, Endo Y, Buckler-White A, Plishka R, Bischofberger N, et al. Macrophage are the principal reservoir and sustain high virus loads in rhesus macaques after the depletion of CD4+ T cells by a highly pathogenic simian immunodeficiency virus/HIV type 1 chimera (SHIV): Implications for HIV-1 infections of humans. Proceedings of the National Academy of Sciences of the United States of America. 2001;98(2):658–63. doi: 10.1073/pnas.98.2.658.

22. Cingöz O, Arnow ND, Puig Torrents M, Bannert N. Vpx enhances innate immune responses independently of SAMHD1 during HIV-1 infection. Retrovirology. 2021;18(1):4. doi: 10.1186/s12977-021-00548-2.

23. Gaidt MM, Ebert TS, Chauhan D, Ramshorn K, Pinci F, Zuber S, et al. The DNA Inflammasome in Human Myeloid Cells Is Initiated by a STING-Cell Death Program Upstream of NLRP3. Cell. 2017;171(5):1110-24.e18. Epub 20171012. doi: 10.1016/j.cell.2017.09.039. PubMed PMID: 29033128; PubMed Central PMCID: PMCPMC5901709.

24. Gao D, Wu J, Wu Y-T, Du F, Aroh C, Yan N, et al. Cyclic GMP-AMP synthase is an innate immune sensor of HIV and other retroviruses. Science. 2013;341(6148):903–6. Epub 08/08. doi: 10.1126/science.1240933. PubMed PMID: 23929945.

25. Johnson JS, Lucas SY, Amon LM, Skelton S, Nazitto R, Carbonetti S, et al. Reshaping of the Dendritic Cell Chromatin Landscape and Interferon Pathways during HIV Infection. Cell Host & Microbe. 2018;23(3):366-81.e9. doi: https://doi.org/10.1016/j.chom.2018.01.012.

26. Rasaiyaah J, Tan CP, Fletcher AJ, Price AJ, Blondeau C, Hilditch L, et al. HIV-1 evades innate immune recognition through specific cofactor recruitment. Nature. 2013;503(7476):402–5. Epub 2013/11/08. doi: 10.1038/nature12769. PubMed PMID: 24196705; PubMed Central PMCID: PMCPMC3928559.

27. Bonifati S, Daly MB, St Gelais C, Kim SH, Hollenbaugh JA, Shepard C, et al. SAMHD1 controls cell cycle status, apoptosis and HIV-1 infection in monocytic THP-1 cells. Virology. 2016;495:92–100. Epub 2016/05/18. doi: 10.1016/j.virol.2016.05.002. PubMed PMID: 27183329; PubMed Central PMCID: PMCPMC4912869.

28. Malim MH, Freimuth WW, Liu J, Boyle TJ, Lyerly HK, Cullen BR, et al. Stable expression of transdominant Rev protein in human T cells inhibits human immunodeficiency virus replication. J Exp Med. 1992;176(4):1197–201. doi: 10.1084/jem.176.4.1197. PubMed PMID: 1402661.

29. Wang Q, Gao H, Clark KM, Mugisha CS, Davis K, Tang JP, et al. CARD8 is an inflammasome sensor for HIV-1 protease activity. Science. 2021;371(6535). Epub 20210204. doi: 10.1126/science.abe1707. PubMed PMID: 33542150; PubMed Central PMCID: PMCPMC8029496.

30. Akiyama H, Miller CM, Ettinger CR, Belkina AC, Snyder-Cappione JE, Gummuluru S. HIV-1 intron-containing RNA expression induces innate immune activation and T cell dysfunction. Nature Communications. 2018;9(1):3450. doi: 10.1038/s41467-018-05899-7.

31. McCauley SM, Kim K, Nowosielska A, Dauphin A, Yurkovetskiy L, Diehl WE, et al. Intron-containing RNA from the HIV-1 provirus activates type I interferon and inflammatory cytokines. Nature Communications. 2018;9(1):5305. doi: 10.1038/s41467-018-07753-2.

32. Sumner RP, Harrison L, Touizer E, Peacock TP, Spencer M, Zuliani-Alvarez L, et al. Disrupting HIV-1 capsid formation causes cGAS sensing of viral DNA. The EMBO journal. 2020;39(20):e103958. Epub 2020/08/28. doi: 10.15252/embj.2019103958. PubMed PMID: 32852081; PubMed Central PMCID: PMCPMC7560218.

33. Gringhuis SI, Hertoghs N, Kaptein TM, Zijlstra-Willems EM, Sarrami-Forooshani R, Sprokholt JK, et al. HIV-1 blocks the signaling adaptor MAVS to evade antiviral host defense after sensing of abortive HIV-1 RNA by the host helicase DDX3. Nature immunology. 2017;18(2):225–35. doi: 10.1038/ni.3647.

34. Monroe KM, Yang Z, Johnson JR, Geng X, Doitsh G, Krogan NJ, et al. IFI16 DNA sensor is required for death of lymphoid CD4 T cells abortively infected with HIV. Science. 2014;343(6169):428–32. Epub 2013/12/21. doi: 10.1126/science.1243640. PubMed PMID: 24356113; PubMed Central PMCID: PMCPMC3976200.

35. Lepelley A, Louis S, Sourisseau M, Law HK, Pothlichet J, Schilte C, et al. Innate sensing of HIV-infected cells. PLoS Pathog. 2011;7(2):e1001284. Epub 20110217. doi: 10.1371/journal.ppat.1001284. PubMed PMID: 21379343; PubMed Central PMCID: PMCPMC3040675.

36. Zhang G, Luk BT, Wei X, Campbell GR, Fang RH, Zhang L, et al. Selective cell death of latently HIV-infected CD4(+) T cells mediated by autosis inducing nanopeptides. Cell Death Dis. 2019;10(6):419. Epub 20190529. doi: 10.1038/s41419-019-1661-7. PubMed PMID: 31142734; PubMed Central PMCID: PMCPMC6541658.

37. Cooper A, García M, Petrovas C, Yamamoto T, Koup RA, Nabel GJ. HIV-1 causes CD4 cell death through DNA-dependent protein kinase during viral integration. Nature. 2013;498(7454):376–9. doi: 10.1038/nature12274.

38. Liu Y, Shoji-Kawata S, Sumpter RM, Jr., Wei Y, Ginet V, Zhang L, et al. Autosis is a Na+,K+-ATPase-regulated form of cell death triggered by autophagy-inducing peptides, starvation, and hypoxia-ischemia. Proceedings of the National Academy of Sciences of the United States of America. 2013;110(51):20364–71. Epub 2013/11/28. doi: 10.1073/pnas.1319661110. PubMed PMID: 24277826; PubMed Central PMCID: PMCPMC3870705.

39. Martin JL, Cao S, Maldonado JO, Zhang W, Mansky LM, Ross SR. Distinct Particle Morphologies Revealed through Comparative Parallel Analyses of Retrovirus-Like Particles. Journal of Virology. 2016;90(18):8074–84. doi: 10.1128/JVI.00666-16.

40. Andersen JL, DeHart JL, Zimmerman ES, Ardon O, Kim B, Jacquot G, et al. HIV-1 Vpr-Induced Apoptosis Is Cell Cycle Dependent and Requires Bax but Not ANT. PLOS Pathogens. 2006;2(12):e127. doi: 10.1371/journal.ppat.0020127.

41. Joshi A, Nyakeriga AM, Ravi R, Garg H. HIV ENV glycoprotein-mediated bystander apoptosis depends on expression of the CCR5 co-receptor at the cell surface and ENV fusogenic activity. The Journal of biological chemistry. 2011;286(42):36404–13. Epub 2011/08/24. doi: 10.1074/jbc.M111.281659. PubMed PMID: 21859712; PubMed Central PMCID: PMCPMC3196103.

42. Chelvanambi S, Bogatcheva NV, Bednorz M, Agarwal S, Maier B, Alves NJ, et al. HIV-Nef Protein Persists in the Lungs of Aviremic Patients with HIV and Induces Endothelial Cell Death. American journal of respiratory cell and molecular biology. 2019;60(3):357–66. Epub 2018/10/16. doi: 10.1165/rcmb.2018-0089OC. PubMed PMID: 30321057; PubMed Central PMCID: PMCPMC6397978.

43. Levin A, Hayouka Z, Friedler A, Loyter A. Over-expression of the HIV-1 Rev promotes death of nondividing eukaryotic cells. Virus Genes. 2010;40(3):341–6. doi: 10.1007/s11262-010-0458-7.

44. Lee S, Karki R, Wang Y, Nguyen LN, Kalathur RC, Kanneganti TD. AIM2 forms a complex with pyrin and ZBP1 to drive PANoptosis and host defence. Nature. 2021;597(7876):415–9. Epub 2021/09/03. doi: 10.1038/s41586-021-03875-8. PubMed PMID: 34471287; PubMed Central PMCID: PMCPMC8603942.

45. Nguyen LN, Kanneganti TD. PANoptosis in Viral Infection: The Missing Puzzle Piece in the Cell Death Field. Journal of molecular biology. 2022;434(4):167249. Epub 2021/09/20. doi: 10.1016/j.jmb.2021.167249. PubMed PMID: 34537233; PubMed Central PMCID: PMCPMC8444475.

46. Dixon SJ, Lemberg KM, Lamprecht MR, Skouta R, Zaitsev EM, Gleason CE, et al. Ferroptosis: an iron-dependent form of nonapoptotic cell death. Cell. 2012;149(5):1060–72. Epub 2012/05/29. doi: 10.1016/j.cell.2012.03.042. PubMed PMID: 22632970; PubMed Central PMCID: PMCPMC3367386.

47. Holze C, Michaudel C, Mackowiak C, Haas DA, Benda C, Hubel P, et al. Oxeiptosis, a ROS-induced caspase-independent apoptosis-like cell-death pathway. Nature immunology. 2018;19(2):130–40. Epub 2017/12/20. doi: 10.1038/s41590-017-0013-y. PubMed PMID: 29255269; PubMed Central PMCID: PMCPMC5786482.

